# Integrated analysis reveals strong reproducible signals within and across studies of the built environment

**DOI:** 10.64898/2026.04.03.716326

**Authors:** Abeoseh B. Flemister, Ivory Blakley, Anthony A. Fodor

**Affiliations:** College of Computing and Informatics, University of North Carolina at Charlotte, 9201 University City Blvd, Charlotte, NC 28223, USA

**Keywords:** Built Environment, Reproducibility, Batch Effects

## Abstract

**Background:** Built environment microbiome studies have identified numerous factors that shape indoor microbiomes, yet the reproducibility of these findings across buildings, timepoints, and research groups remains unclear. Differences in sequencing protocols, sampling design, and environments pose major challenges for cross-study comparisons, particularly in low-biomass environments where technical variation can obscure biological signal. To address this gap, we constructed a simple ontology which groups samples into one of three categories: hand, hand-associated surfaces, and floor then applied it to four publicly available 16S rRNA gene datasets: a hospital, university dormitory, Air Force dormitory, and private residential houses.

**Results:** We identified strong and reproducible separation between floors and surfaces with frequent human contact. We found that floors consistently harbored soil-associated taxa, including *KD4-96, 67-14, Skermanella,* and *Sphingobacterium*, whereas hands and hand-associated surfaces were enriched with skin-associated genera, such as *Lawsonella* and *Cutibacterium*. Within studies, these results were generally consistent across timepoints. Across studies, mixed-model PERMANOVA analysis revealed significant clustering by sample type, with modest effects of study, suggesting that biological signal outweighed differences in laboratory or sequencing methods. Leave-one-study-out random forest models achieved high AUCs for hand vs. floor comparisons (0.865 to 0.921), moderate AUCs for hand-associated vs. floor comparisons, and weaker performance for hand vs. hand-associated comparisons. Application of the batch-correction method DEBIAS-M did not improve effect sizes or classification performance, indicating that reproducible structure was already discernible without batch adjustment.

**Conclusions:** Despite substantial temporal and environmental heterogeneity among studies, we found that the built environment microbiome has a reproducible bacterial signal. There was consistent enrichment of soil-derived taxa on floors and human-associated taxa on hands and hand-associated surfaces suggesting a stable microbiome despite differences in building type, occupancy, and methodology. These findings establish an important foundation for future studies, suggesting cross-study comparability, the accuracy of ecological inference, and the ability to support the development of predictive applications in indoor microbiome research.

## Background

The built environment comprises man-made structures which are distinct from the natural environment. Within these indoor spaces, microorganisms are abundant and can significantly impact human health, promoting both wellness and disease. The complex bacterial communities of the built environment are influenced by multiple factors including geographical location^1^, the characteristics and behaviors of human occupants^2^, and the time of year^3^. Previous studies have described temporal variability within the built environment^3–5^ and colonization events within newly opened or cleaned buildings^4–6^. Studies have also described seasonal variation in the microbiome of buildings^3^ with humans activity acting as vectors for microbes.^7,8^ While individual studies produce valuable insights into what determines microbiome structure, the reproducibility of these observations across studies is uncharacterized.

There are numerous factors which can potentially drive similarities and differences across studies. Microbiome differences in human populations in different geographical regions could cause different studies to detect distinct microbiomes^9^. Conversely, consistent selection pressures on different regions of the built environment, such as skin-derived microbes populating high-touch areas and soil-derived microbes accumulating on floors, could cause convergence across studies despite differences in occupants or usage patterns^10^. Additionally, reproducibility in microbiome studies could also be impacted by differences in technical and analytical methods, including sample collection protocols, type of extraction kit, PCR and sequencing platform used, as well as bioinformatics pipelines.^11–14^ These challenges are significant for any microbiome study but become more pronounced in low-biomass environments, such as the built environment, where contamination and technical noise can overwhelm biological signal.^15,16^

Technical variation can produce strong batch effects in microbiome data that have the potential to overwhelm real signals, inflate false-positive rates, reduce statistical power, and create misleading ecological inferences. Currently, several tools are available to correct for batch correction including ComBat^17^, ConQuR^18^, percnorm^19^, Voom-SNM^20,21^, MMUPHin^22^ and PLSDA-batch^23^. These algorithms differ in their assumptions and performance across data types. However, batch-correction methods developed for high-biomass datasets or gene-expression data may not generalize well to microbiome studies, particularly when the data is sparse, compositional, or zero-inflated. In this study, we focused on DEBIAS-M, a recently proposed algorithm designed specifically to address batch effects in microbiome datasets by modeling and removing study-specific biases without distorting underlying ecological relationships.

To address these issues, we compare four publicly available 16S rRNA gene datasets. These studies represent four disparate environments (hospital^4^, university dormitory^24^, Air Force dormitory^3^, and households^25^). To compare studies with distinct yet complementary naming conventions and sample types, we created a simple ontology to group samples into hand, hand-associated surfaces, and floor. Utilizing multivariate statistics, pairwise statistical inference, and leave-one-study-out random forest we found a surprising degree of agreement across studies in separating hands from floors and floors from hand-associated surfaces, but a much less robust signal when comparing hands with hand-associated surfaces. Our study suggests a common partitioning of the built environment that yields a reproducible signal across studies, despite differences in the environment sampled and methodological difference in sampling.

## Methods

### Dataset Selection

To examine patterns of reproducibility across built environment studies, we searched NCBI’s SRA and BioProject databases for “text word” or “any field” containing “hand”, “skin”, “floor”, and any surfaces associated with hands. Additionally, we searched NCBI PubMed and Google Scholar for studies with publicly available datasets that had 16S rRNA gene bacterial sequences in FASTQ format with sample metadata that fit within our ontology. All studies were sequenced on Illumina platforms. We found four studies (**Table 1; Table S1**) which met our criteria studies within: A hospital, university dormitories, Air Force dormitories, and houses. Table 1 shows our sample breakdown before and after any sub-sampling occurred. Lax et al. 2017: hospital had 3079 of the 6523 reported samples in the SRA; therefore, we were unable to utilize the full dataset. For each comparison among studies, we created an ontology to group related surfaces (**Table 1**).

**Table 1:** Study characteristics. The total amount of each sample (Total Samples), the amount used (Used Samples), the name used in the study (Name), and the common name we used in our analysis (Common Name) for the hand, hand associated, and floor samples for each study (Dataset). Sharma et al. 2019: Air Force swabbed the inner elbow.

| Dataset | Total Samples | Used Samples | Name | Common Name |
| --- | --- | --- | --- | --- |
| Lax et al. 2017: Hospital | 304 | 286 | Hand | Hand |
|  | 477 | 455 | Floor (414), Corridor Floor (41) | Floor |
|  | 1026 | 985 | cold tap faucet handle (243), bedrail (257), countertop (43), computer mouse (42), station phone (42), chair armrest (43), personal cell phone (137), hospital pager (139), hot tap water faucet handle (20), cold tap water faucet handle (19) | hand associated surfaces |
| Sharma et al. 2019: Air Force | 234 | 229 | skin | Hand |
|  | 93 | 86 | floor corner (37), squad common room (49) | Floor |
|  | 535 | 535 | Desk (495), bathroom handle (40) | hand associated surfaces |
| Richardson et al. 2019: Dorm | 114 | 114 | Hand | Hand |
|  | 93 | 93 | Floor (80), hallway (13) | Floor |
|  | 224 | 224 | Desk (79), door handle (78), elevator button (18), main door (13), table (16), bathroom (20) | hand associated surfaces |
| Lax et al. 2014: House | 279 | 279 | Hand | Hand |
|  | 229 | 229 | bedroom floor (115), kitchen floor (114) | Floor |
|  | 460 | 457 | Bathroom doorknob (114), front doorknob (115), kitchen counter (114), kitchen light switch (114), | hand associated surfaces |

### Ontology to determine reproducibility in built environment datasets

Our analysis asks how hand, hand-associated, and floor environments differ from each other across time and across studies. For our four publicly available datasets (**Table 1**), we created an ontology assigning samples in these datasets to hand, hand-associated, and floor (**Table 1 and S1**). Hand-associated surfaces represent environmental samples that are in contact with human hands. Three of the four studies sampled human hand and these were identified in the study as “Hand”. Dorm: Sharma et al., 2019, used the inner elbow as their hand sample which we also grouped under hand in our ontology. Each study had distinct names for hand-associated surfaces (such as desk, countertop, or bedrail) and using our ontology and we collected all of these terms as “hand associated” (**Table 1**). Likewise, the studies had many different names for floor samples (such as “floor” or “floor corner” or “bedroom floor”), for most of these samples the word “floor” was usually present. We collected all these terms under the word floor in our simple ontology.

### Timepoint and Longitudinal Analyses

For the timepoint analysis, we selected timepoints which had at least 30 samples for hand, hand-associated or floor. For pairwise comparisons (i.e., hand vs floor, hand-associated vs floor, and hand vs hand-associated) we only used timepoints which had 30 samples for both ontological groups (**Table S1**). This led us to only use the hospital study for the hand vs floor comparison (three timepoints), the hospital and dorm studies for the hand-associated vs floor comparison (six and two timepoints respectively) and the hospital and Air Force studies for the hand vs hand-associated comparison (four timepoints each). Since all the ontological groups had at least 30 samples for the longitudinal comparisons, we did not employ similar filtering techniques, and we used all studies for all comparisons (**Table 1**).

### Taxonomic Assignment

We performed denoising with QIIME 2 2024.10^26^ DADA2^27^ denoise-single with no trimming or truncation. We pooled and removed chimeras. Sequences were considered chimeras if the parent sequences were 100-fold more abundant. Taxonomic assignment was done using Qiime2 feature-classifier classify-sklearn^28^ which is a naive bayes kmer classifier trained on the experimental human stool weighted Silva 138 99% OTUs full-length sequences database^28–30^.

### Log Transformation

To correct for read depth prior to analysis, we performed a log-normalization, as described in Yerke et al., 2024^31^.

### Statistical Methods and Analyses

#### PERMANOVA and PCoA

All statistical analyses were done in an R_v4.3.3_ environment^32^. We conducted PERMANOVA using adonis2 from the vegan package ^33^ with 100000 or 1000 permutations. PCoA was calculated using the ecodist package ^34^ based on Bray-Curtis dissimilarity metric.

#### Relative abundance plots

For the timepoint comparisons, we created relative abundance plots using ggplot2_v3.5.1_^35^ to identify the proportion of each bacterium in each sample The plots were created in R and visualized the top 20 most abundant bacteria with the remaining bacteria labelled as other.

#### p-value vs p-value plots

For each bacterium within each study, we calculated a Wilcoxon rank sum test between our pairwise groups (i.e., hand vs floor, hand-associated vs floor, and hand vs hand-associated). When the pairwise comparison included floor samples, we made the log10 p-value positive if the mean of the hand or hand-associated samples was greater than the mean of the floor samples. For the hand vs hand-associated comparison, the log10 p-value was weighted as positive if the mean for the hand samples was greater than the mean of the hand-associated samples. We then plotted all the bacteria present in both the studies against each other where the first quadrant shows agreement between studies that a given bacterium was present in higher quantities in the positively weighted samples while the third quadrant shows agreement between studies that a given bacterium was present in higher quantities in negatively weighted samples.

### Random Forest

#### Timepoint analyses

To compare microbiome changes across time, for each study with timepoints that had at least 30 samples, we did Random Forest using the randomForest package ^36^. The AUC, sensitivities, and specificities were calculated using the pROC_v1.18.5_ package^37^. To determine the statistical significance of our model’s ability to predict differences in the microbiome between surface types, we performed 100 permutations where we permuted the labels for both training and testing datasets. We then calculated a permutation p-value as the proportion of the permutated AUCs greater than or equal to the un-permutated AUC. When training and testing on the same timepoint, we performed 10-fold cross validation with a 70-30 training-testing split and computed the average AUC.

#### Longitudinal analyses

To elucidate reproducibility across study, we performed Random Forest using the randomForest package _v4.7-1.1_^36^ in R using a leave-one-study-out (LOSO) design. For each pairwise comparison, we trained on three out of four datasets and tested on the fourth. To determine the relative ability of our model to separate the two classes, we performed 1000 permutations where both the training and testing labels were randomized. The AUC, sensitivities, and specificities were calculated using the pROC package^37^. We then calculated a permutation p-value as the proportion of the permutated AUCs greater than or equal to the un-permutated AUC.

## Results

### Ontology

Our study included four publicly available datasets that had sufficient sample size to perform machine learning. A central challenge in comparing across studies is that different studies used different terminology to define their samples. To address this, we created a simple ontology where we translated the terms used in each study into grouped samples that we designated as hand, hand-associated, and floor. **Table 1** shows our ontology applied to all samples in our four studies while **Table S1** shows samples broken out by timepoint.

### Timepoint Analysis Reveals Disparate Microbiomes between Hands and Floors

Our analysis begins with an examination of the reproducibility across timepoints **within** each study. We will then next address reproducibility **across** studies.

We begin with the hospital study, which is the largest of all the studies in our datasets with over 1,500 samples spread out over seven timepoints. First, we examine the comparison where we anticipated the greatest difference between microbiome composition, floor vs. hand. Of the seven timepoints in the hospital study, only three (2013/03, 2013/07, 2013/12) had at least 30 samples for both hand and floor samples **(Table S1)**. We set 30 samples as the minimum number of samples that could reasonably power machine leaning models. Visualization of the 20 most abundant bacteria from the hands and floor samples from these timepoints shows a very clear separation between hands and floors **(Fig. 1A)**. There was a strong separation between hands and floors for two of the three timepoints, with a much weaker separation for the July 2013 timepoint (**Fig. 1B**).

**Figure 1:**
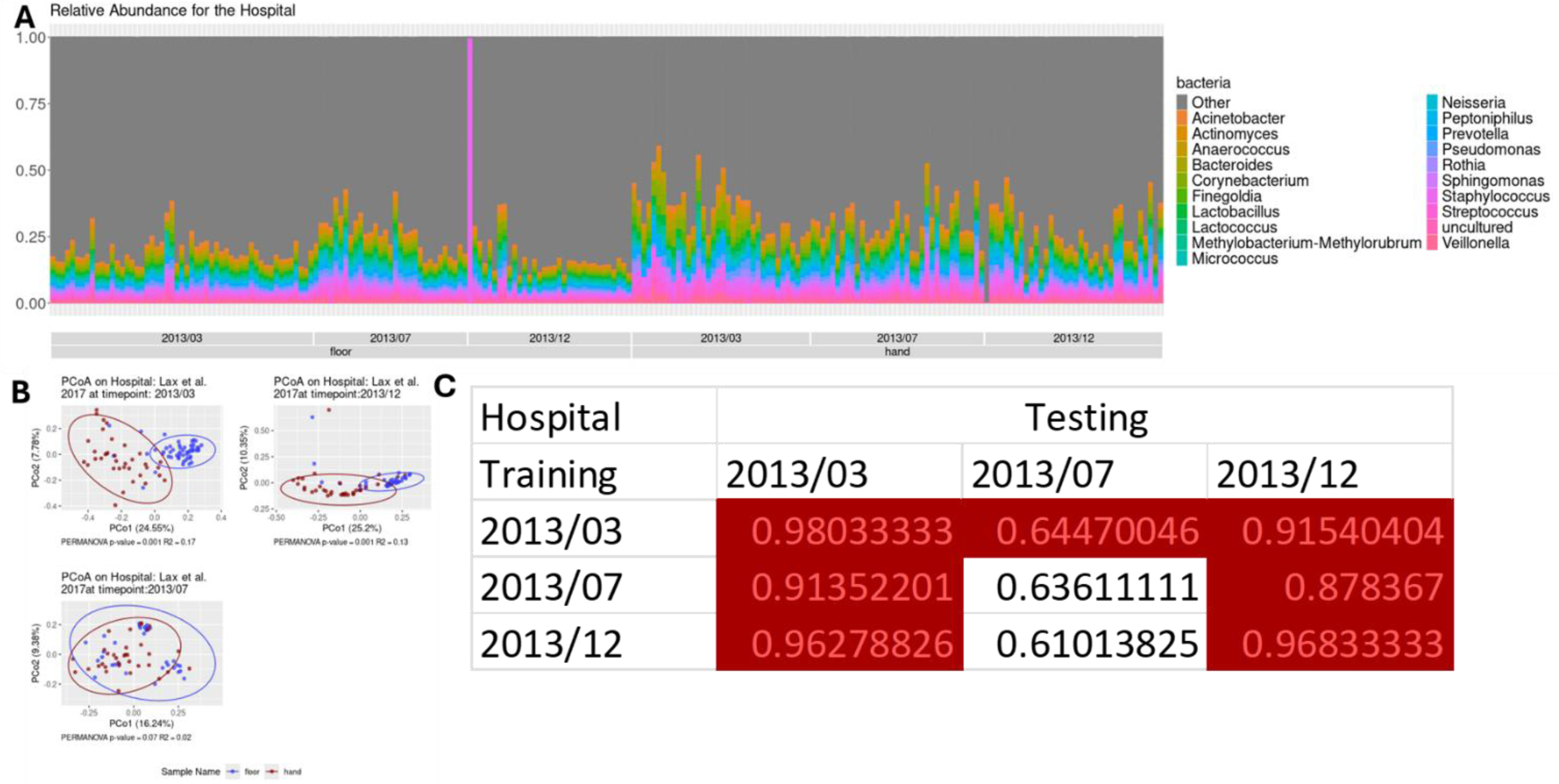
How the microbiome of hands and floors change over time within the Hospital study. **(A)** Relative abundance between hands and floors for the top 20 bacteria and others, **(B)** PCoAs and PERMANOVA with 1,000 permutations between timepoints 2013/03, 2013/12, and 2013/07. **(C)** AUC for pairwise random forests with 100 permutations. The AUC is for the un-permutated data and the p-value (significance denoted by a red box) is the number of times the AUC of the permutations were equal to or greater than the AUC of the actual data.

We trained random forest models on one timepoint and tested on the others individually. In general, training on all three timepoints created models that could be successfully used to make predictions on the other two timepoints; however, when using the timepoints for inference, the 2013/07 timepoint generated substantially lower AUCs. We note that the 2013/07 PCoA lacks separation between hand and floor samples (**Fig. 1B**). We speculate that the random forest algorithm was able to learn intricate patterns in 2013/07 which allows it to predict the much broader differences in the other timepoints. But since the hand and floor are heterogeneous for this timepoint, the models from the other dates cannot reliably predict differences for this date.

The hospital study was the only study with more than 30 samples for both hand and floor samples across multiple timepoints, and we, therefore, do not consider hand vs. floor comparisons within the other three studies.

### Comparison between Hand-Associated Surfaces and Floors reveals strong separation

The Hospital and Dorm studies had six and two timepoints, respectively, which had at least 30 samples for hand-associated and floor samples. Unlike the floor vs. hand comparison, hand-associated surfaces and floors are both environmental; we therefore expected less separation than between hand and floor samples. However, most of the time points for the hospital study **(Fig. 2)** and all the time points for the dorm study **(Fig. 3)** showed very clear separation between floors and hand-associated surfaces. PCoAs for the hospital study **(Fig. 2B)** showed significant separation for all time points, however, the 2013/07 timepoint showed less separation than the other timepoints. Our time point random forest models were capable of predicting the separation between hand-associated surfaces and floors on all time points except 2013/07 **(Fig. 2C)**.

**Figure 2:**
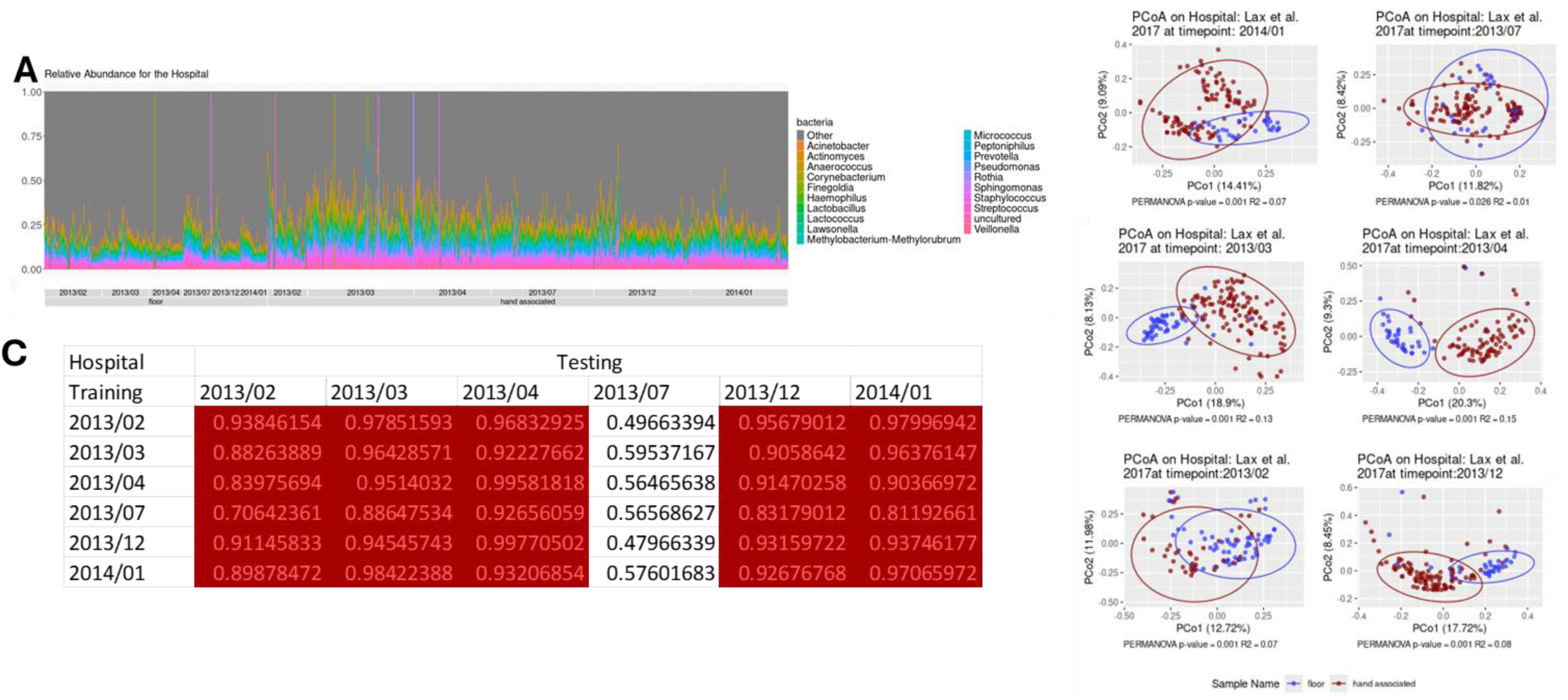
How the microbiome of hand associated surfaces and floors change over time within the Hospital study. **(A)** Relative abundance between hand associated surfaces and floors for the top 20 bacteria and others, **(B)** PCoAs and PERMANOVA with 1,000 permutations between timepoints 2013/02, 2013/03, 2013/04, 2013/07, 2013/12, and 2014/01. **(C)** AUC for pairwise random forests with 100 permutations. The AUC is for the un-permutated data, and the p-value (significance denoted by a red box) is the number of times the AUC of the permutations were equal to or greater than the AUC of the actual data.

**Figure 3:**
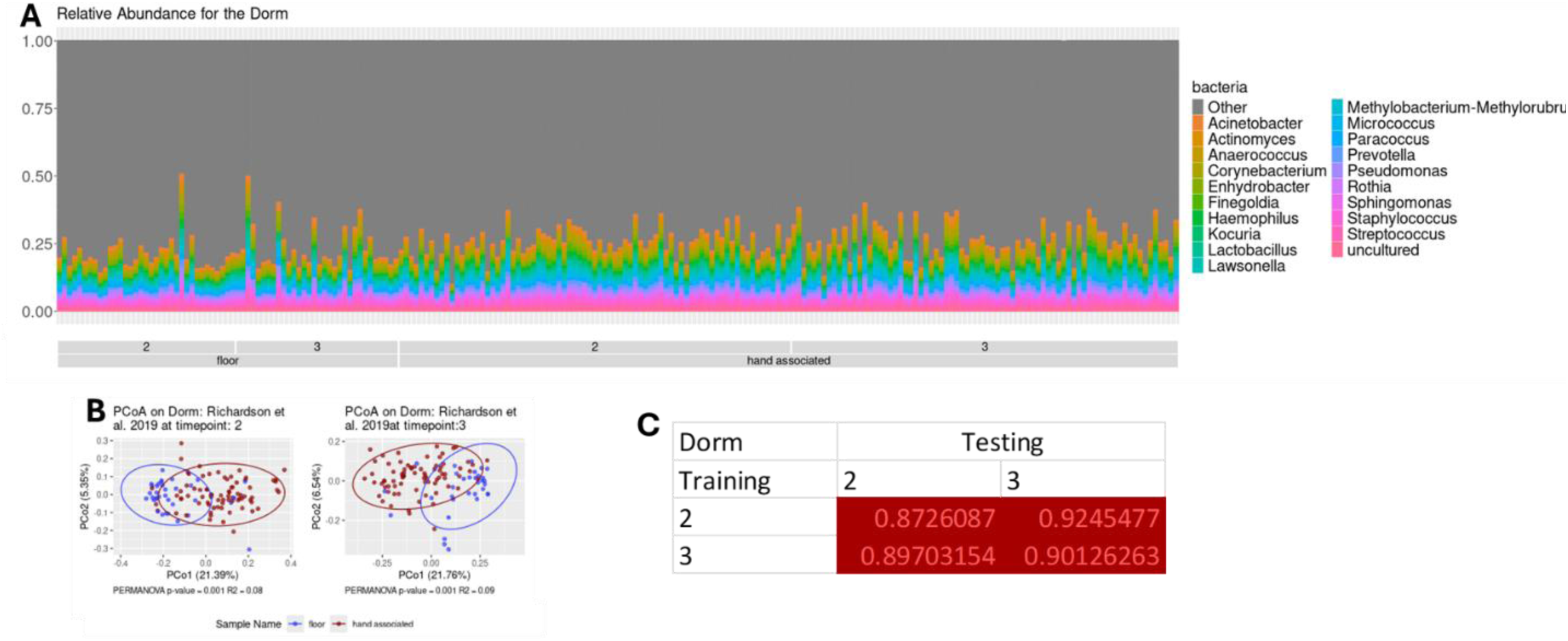
How the microbiome of hand associated surfaces and floors change over time within the Dorm study. **(A)** Relative abundance between hand associated surfaces and floors for the top 20 bacteria and others, **(B)** PCoAs and PERMANOVA with 1,000 permutations between timepoints 2 and 3. **(C)** AUC for pairwise random forests with 100 permutations. The AUC is for the un-permutated data and the p-value (significance denoted by a red box) is the number of times the AUC of the permutations were equal to or greater than the AUC of the actual data.

Mirroring the hospital study, the two time periods within the Dorm study had stable microbiomes within hand-associated surfaces and floor and clear separation between these ontological groups **(Fig. 3A&B)**. Our random forest timepoint models had strong predictive ability across and within timepoints **(Fig. 3C)**.

### Timepoint analysis revealed a less consistent signal in comparing Hand-Associated surfaces and Hands

To conclude our within study comparisons, we looked at the hand vs. hand-associated samples. Since hand and hand-associated are close in physical space, we might anticipate broad similarity between these two environments.

The Hospital and Air Force studies each had four timepoints which had at least 30 samples for hand-associated surfaces and hands. The hospital study displayed similarity in the microbiome of hand-associated surfaces and hands **(Fig. 4A&B)**. Only the 2013/03 (PERMANOVA p-value = 0.003; **Fig. 4B**) and May 2013 (PERMANOVA p-value = 0.031; **Fig. 4B**) timepoints were significantly different but the degree of separation in even these timepoints was modest. These results were mirrored in the ROC curves, with few pairwise ROC curves showing significant separation from their permutations **(Fig. 4C)**.

**Figure 4:**
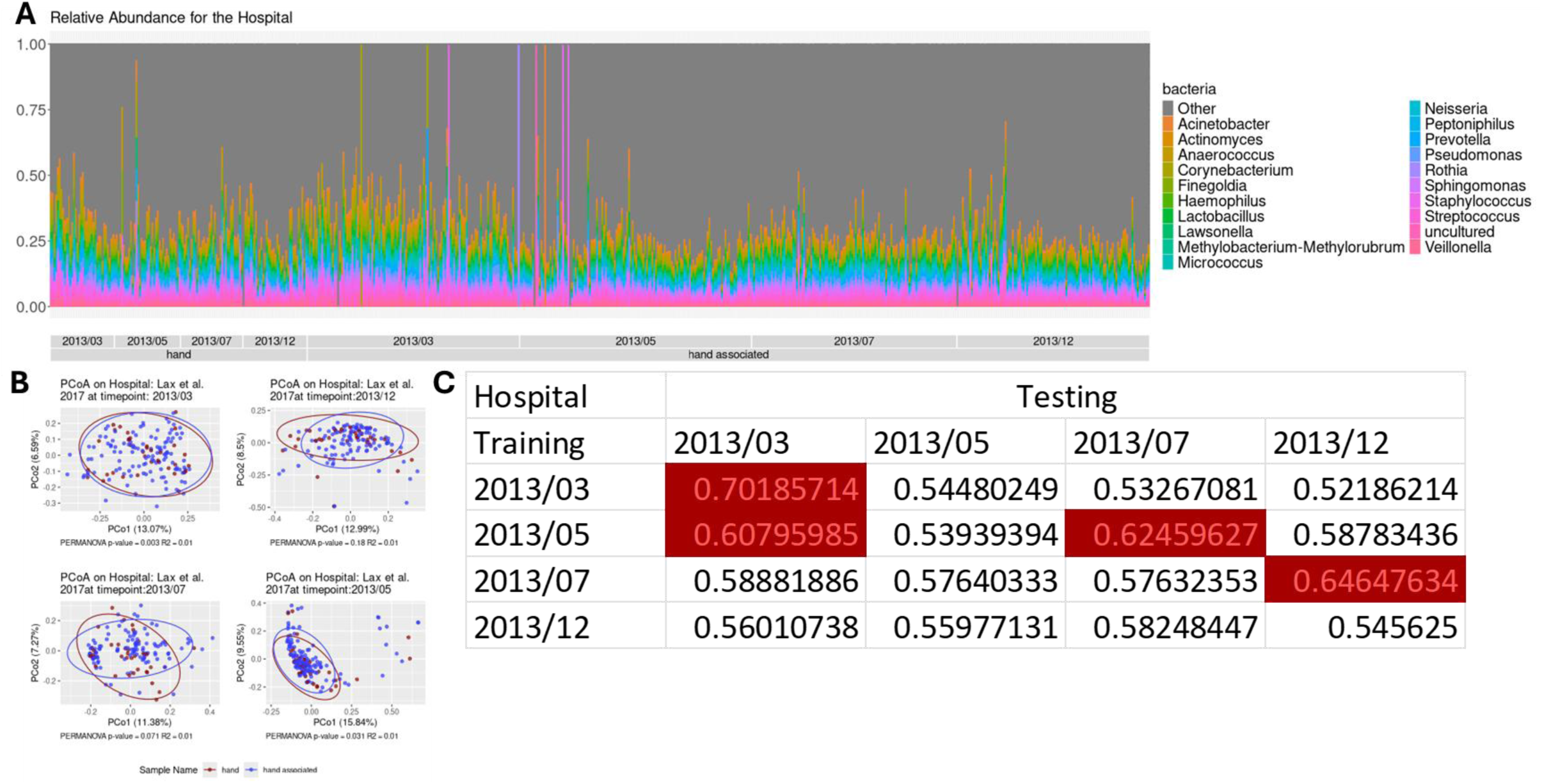
How the microbiomes of hands and hand associated surfaces change over time within the Hospital study. **(A)** Relative abundance between hand associated surfaces and floors for the top 20 bacteria and others, **(B)** PCoAs and PERMANOVA with 1,000 permutations between timepoints 2 and 3. **(C)** AUC for pairwise random forests with 100 permutations. The AUC is for the un-permutated data and the p-value (significance denoted by a red box) is number of times the AUC of the permutations were equal to or greater than the AUC of the actual data.

By contrast, the Air Force study showed strong separation between hand-associated surfaces and hands across all 4 timepoints (**Fig. 5**). The relative abundance plot shows there was large variation with an expansion of *Streptococcus* and *Corynebacterium* within the hand sample types **(Fig. 5A)**. There was likewise pronounced separation in the PERMANOVA plots at all timepoints **(Fig. 5B)**. These results were mirrored in the ROC curves with showed a strong predictive ability across and within each timepoint **(Fig. 5C)**. As noted in the discussion, we don’t have an explanation for the differences between the modest differences in hand vs. hand-associated for the hospital and the strong differences between the dorms, but it is not difficult to imagine hypotheses involving differences in cleaning regiments or resident dwell times in a hospital vs. a university dorm environment.

**Figure 5:**
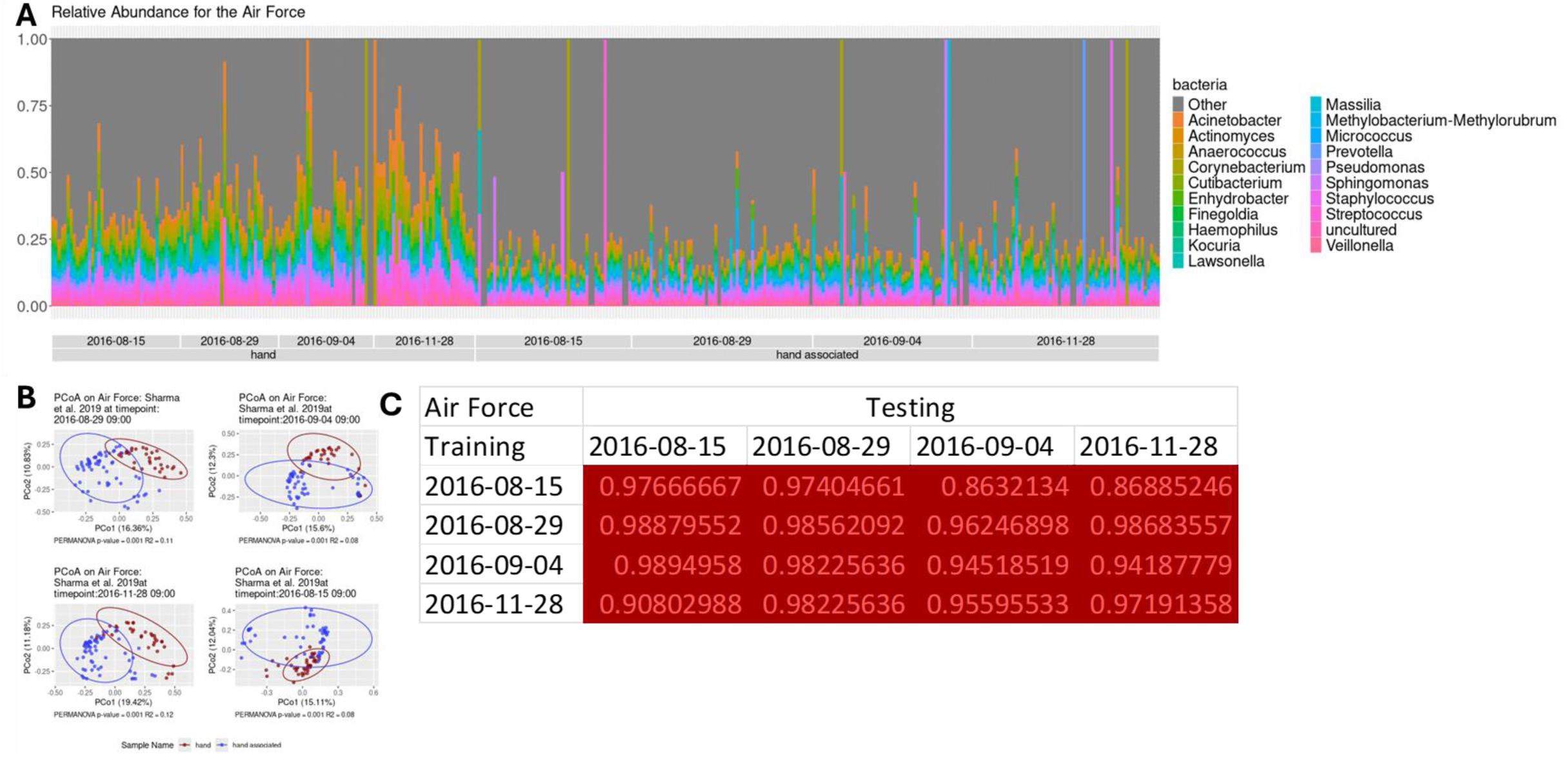
How the microbiomes of hands and hand associated surfaces change over time within the Air Force study. **(A)** Relative abundance between hand and hand associated surfaces for the top 20 bacteria and others, **(B)** PCoAs and PERMANOVA with 1,000 permutations between timepoints 2016-08-15 09:00, 2016-08-29 09:00, 2016-09-04 09:00, and 2016-11-28 09:00. **(C)** AUC for pairwise random forests with 100 permutations. The AUC is for the un-permutated data and the p-value (significance denoted by a red box) is amount of times the AUC of the permutations were equal to or greater than the AUC of the actual data.

### Comparing hand and hand-associated samples across studies revealed a significant but small effect size for clustering by study

Having established the degree of reproducibility within each timepoint, we next examined the degree of reproducibility across studies.

Our studies were from different research institutions with different sequencing and extraction protocols; we therefore begin our analysis by examining how strongly the microbiome of each study clusters. We expected to find a strong study effect in our built environment datasets, however, while there was significant clustering by study, the degree of separation by study was modest with three of the studies largely overlapping in distribution in the PCoA **(Fig. 6A-6C)**. PERMANOVA analysis with study as the univariate variable found significant clustering for all studies but with a low R^2^ (0.04 for all samples, 0.03 for hand samples, and 0.05 for environmental samples) suggesting much of the variation within the data was not explained by study **(Fig. 6)**. T-tests performed on each PCoA axis (**Fig. 6D and 6E**) confirmed that clustering by study was significant but that there was generally substantial overlap between studies.

**Figure 6:**
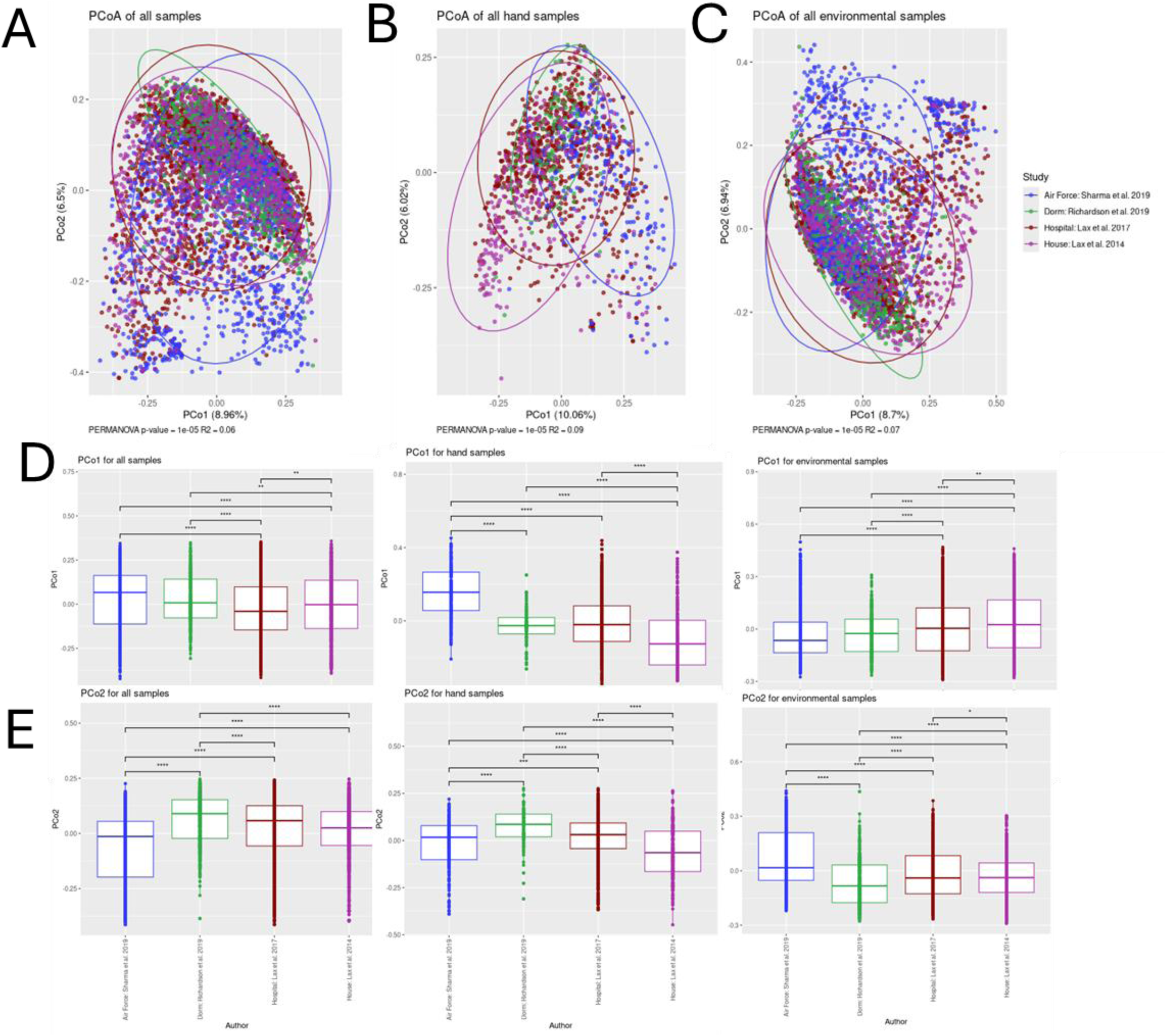
Microbiome differences across study. PCoA based on Bray-Curtis clustered by study for **(A)** all samples, **(B)** hand samples, and **(C)** environmental samples. **(D)** Plot of the first Principal component with a t-test between studies for all samples, hand samples, and environmental samples. **(E)** Plot of the second Principal component with a t-test between studies for all samples, hand samples, and environmental samples.

### Comparisons between hands and floors revealed distinct microbiomes

We next compared hand versus floor samples across studies. The hand vs. floor signal is highly consistent across the studies of these very different built environment habitats. PERMANOVA analysis on all the data with hand versus floor as the fixed term and study as the random term yielded a highly significant PERMANOVA p-value of 1*10^-5^ with an R^2^ value of 0.06 **(Fig. 7A)**. When separated by study, all four studies were significant when comparing hand and floor (p-value < 1*10^-5^) with R^2^ values ranging from 0.07 to 0.14. Despite using the inner elbow, the Air Force study performed just as well as the other studies with an R^2^ of 0.09, only second to the Dorm study with an R^2^ of 0.15 **(Fig 7C&D).**

**Figure 7:**
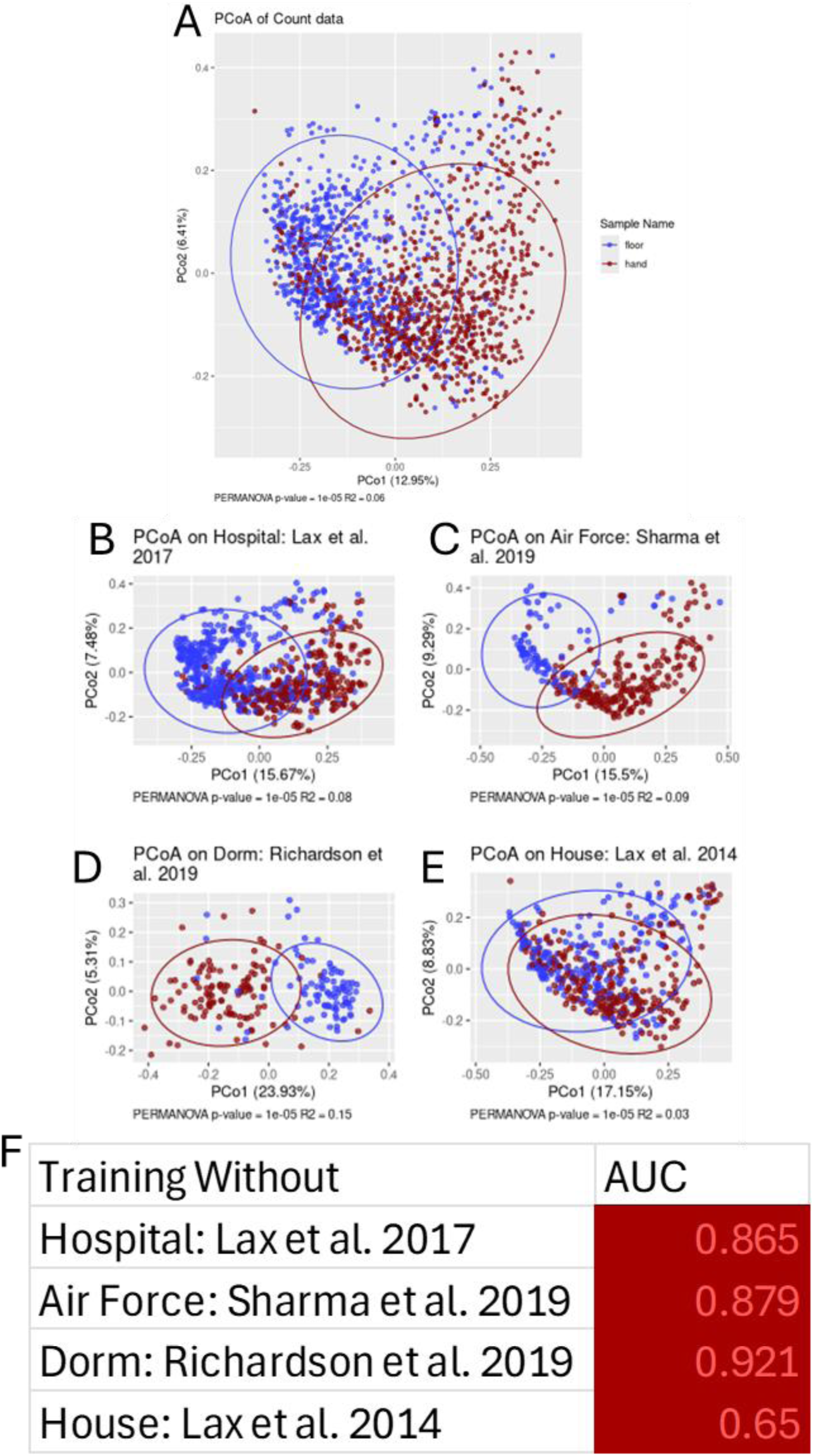
Across study microbiome comparison of hands and floors. PCoA based on Bray-Curtis clustered by **(A)** hand and floor. Hand versus floor PCoA for **(B)** Hospital: Lax et al. 2017, **(C)** air force: Sharma et al. 2019 **(D)** Dorm: Richardson et al. 2019, and **(E)** House: Lax et al. 2014. **(F)** Table of AUC for the unpermuted data in a LOSO design. A red box indicated the p-value was significant for the left out study.

We next took a machine learning approach to ask how reproducible the hand vs. floor signal was across studies. For all the studies, the ROC curve of the un-permutated data cleanly separated **(Fig. S2)** from the ROC curves of the permutations **(Fig 7F)**. The Dorm study complete separation from its permutations with an AUC of 0.921, p < 0.001; the hospital (AUC: 0.865, p < 0.001), Air Force (AUC: 0.874; p-value<0.001), and house studies (AUC: 0.65; p<0.001) followed the same pattern **(Fig. 7F).**

We then sought to determine whether DEBIAS-M, an experimental batch correction algorithm, made an appreciable impact on our observed batch effects and random forest results. A random forest post batch correction with DEBIAS-M did not reveal consistent improvements for AUC the hand and floor comparison **(Fig. S3A&B)**. Permutation p-values did not significantly differ before and after batch correction **(Fig. S3C)**. When performing a PCOA and PERMANOVA with 100,000 permutations, we still found significant clustering by study **(Fig. S3D)** suggesting DEBIAS-M also did not eliminate batch effects.

To elucidate which taxa are responsible for the distinction between hands and floors across study, we created p-value-p-value plots which display the results of inference from a pair of studies **(Fig S4)**. Since we had four datasets there are six possible pairwise comparisons that can be visualized in p-value vs. p-value plots **(Fig. S4)**. Soil bacteria *67-14, KD4-96,* and *Skermanella, Alkanindiges,* and *Sphingobacterium* alongside W*issella*, a pathogenic bacterium, were all consistently significantly more prevalent on floors for multiple studies **(Fig. S4)**. More bacteria were mutually more prevalent in floor samples than hand samples. Comparisons with the house study, which had the smallest size, yielded fewer significant bacteria, and less agreement with the other studies **(Fig. S4D-F)**.

### Comparisons between hand-associated surfaces and floors revealed a less distinct microbiome associated with these environments

We next compared the microbiome of hand-associated surfaces to floor across all studies. The mixed-model PERMANOVA comparing the microbiome of hand-associated surfaces to floor was significant for all comparisons **(Fig. 8A-E)**. When taking a machine learning approach, all studies were significantly able to separate hand-associated surfaces from floors except the Air Force Study which showed little evidence of separation (AUC: 0.506; p=0.769; **Fig. 8F**). We again did not observe an appreciable improvement in hand-associated vs. floor assignments with DEBIAS-M batch correction **(Fig. 8F & Fig. S5)**.

**Figure 8:**
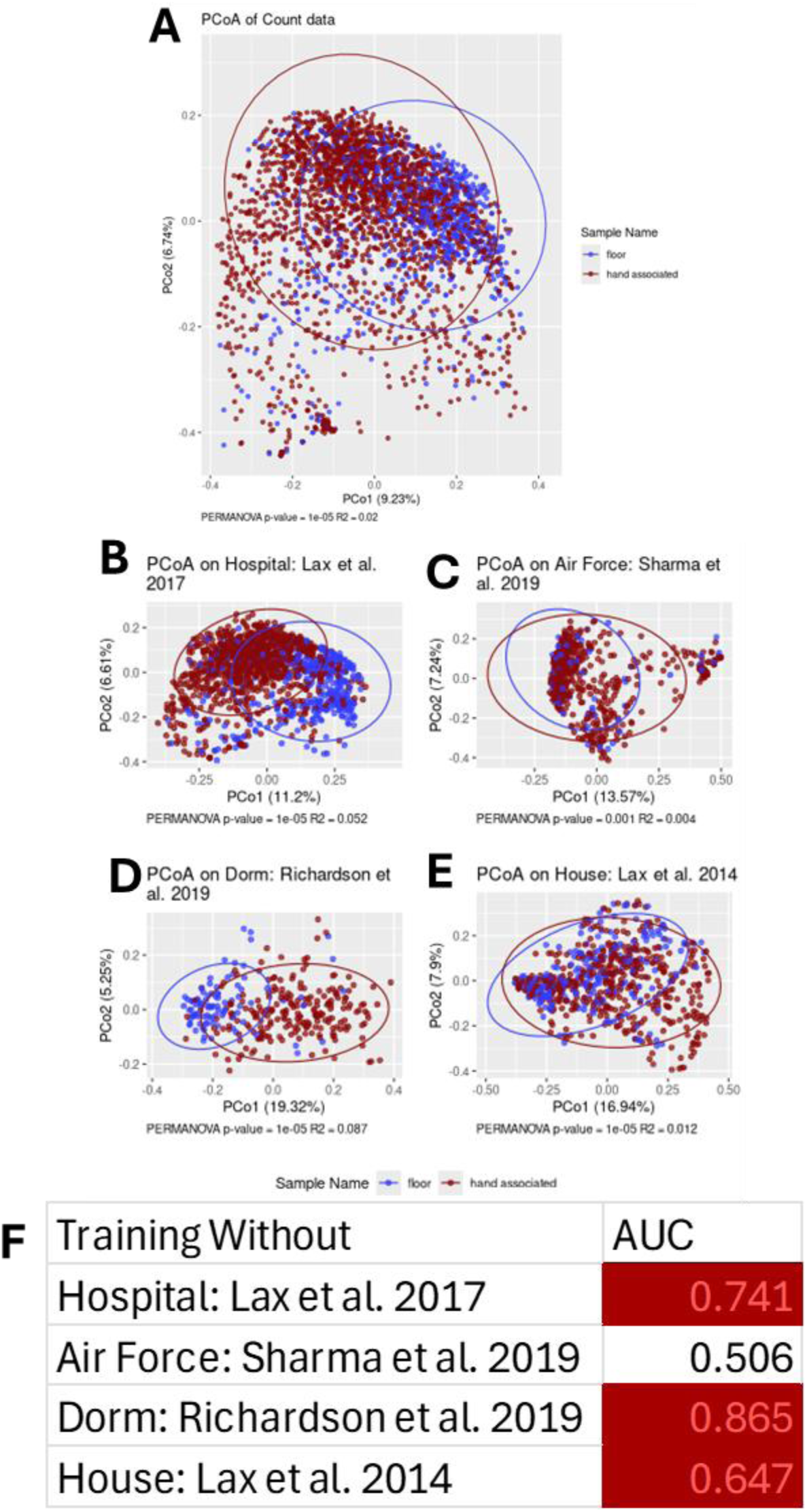
Across study microbiome comparison of hand associated surfaces and floors. PCoA for all samples based on Bray-Curtis distance clustered by **(A)** hand associated and floor. Hand associated versus floor PCoA for **(B)** Hospital: Lax et al. 2017, **(C)** air force: Sharma et al. 2019 **(D)** Dorm: Richardson et al. 2019, and **(E)** House: Lax et al. 2014. **(F)** Table of AUC for the unpermuted data in a LOSO design. A red box indicated the p-value was significant for the left out study.

Similar to our hand versus floor comparisons, our log_10_ p-value-p-value plots indicated that multiple soil bacteria such as *Skermanella, 67-14* and *KD4-96* were mutually present among studies in higher concentrations on floors while human associated bacteria such as *Streptococcus, Lawsonella,* and *Cutibacterium* were mutually present among studies in higher concentrations on surfaces which commonly interact with hands **(Fig. S6A-D)**. Comparisons between the hospital and dorm study revealed strong agreement concerning which bacteria were significantly more prevalent in hand-associated surfaces or floor samples **(Fig. S6A)**. Comparisons between other studies were weaker, with quadrant one (which shows agreement between studies concerning which samples were significantly more prevalent in hand samples) being underpopulated or unpopulated **(Fig. S6B-F)**. This was particularly true for comparisons for the Air Force study **(Fig. S6D-F)**.

### Comparisons between hands and hand-associated surfaces revealed nearly convergent microbiomes

Our final comparison across studies compared hand and hand-associated surface samples. As we might expect, this comparison yields a weaker signal when compared to hand vs. floor and hand-associated vs. floor.

Our mixed-model PERMANOVA revealed significant clustering of samples (p-value < 1x10^-5^) with low variance explained by the model (R^2^=0.01) when collapsing data across all the studies **(Fig 9A)**. Breaking out each study individually, all four studies showed significant differences between hand and hand-associated surfaces, but three of the studies showed a smaller effect size (with r-squared values <=0.02) while the Air Force study (**Fig. 9B-E**) showed a much more pronounced difference (with an r-squared value of 0.08). The results from the leave one study out random forest models were modest with ROCs ranging from 0.54 to 0.69 suggesting a limited ability to distinguish between hand and hand-associated surfaces **(Fig. 9F)**. For three of the four studies, DEBIAS-M made the random forest performance worse, but treatment with DEBIAS-M did lead to an improvement in the Air Force study **(Fig. S8)**.

**Figure 9:**
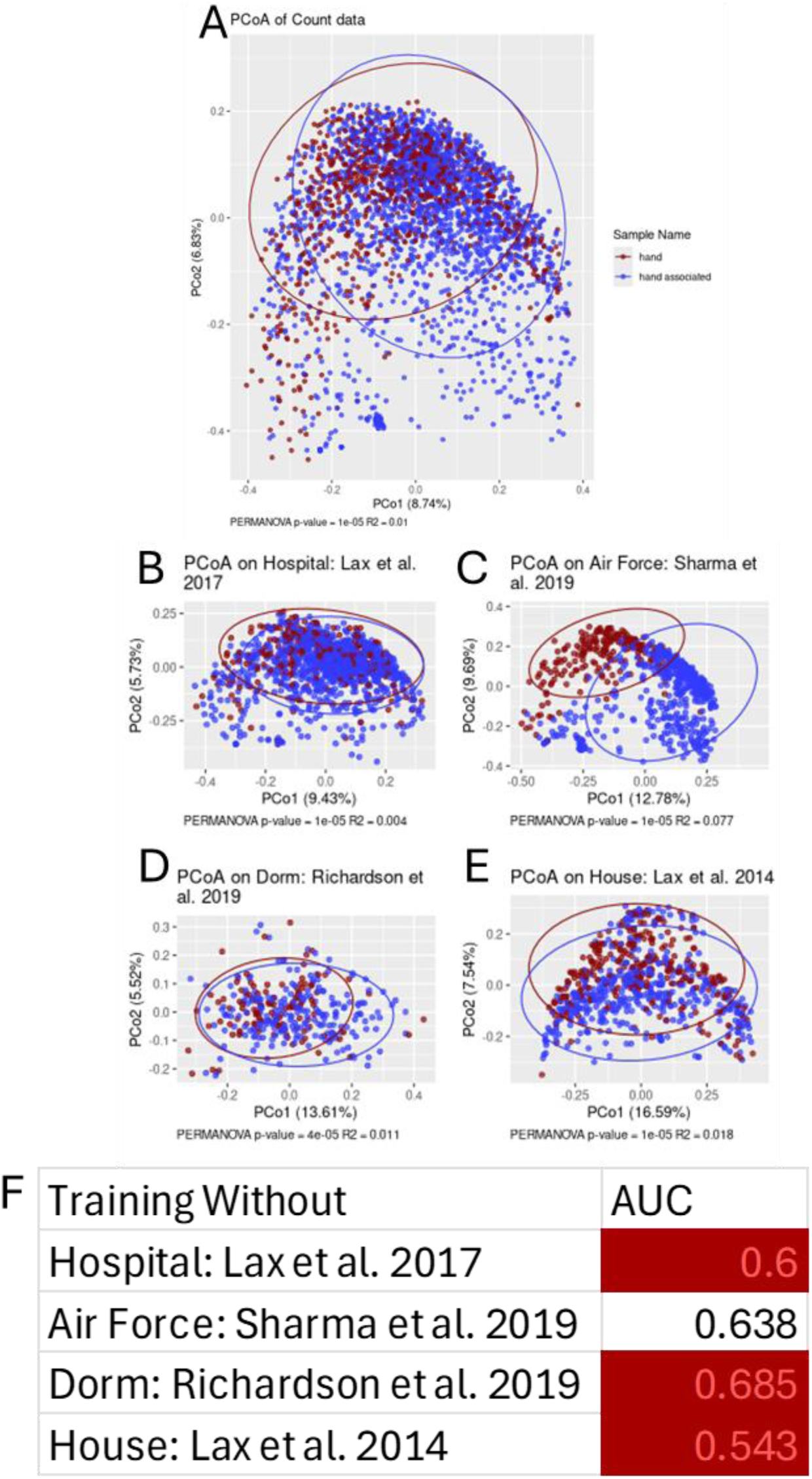
Across study microbiome comparison of hands and hand associated surfaces. PCoA based on Bray-Curtis distance plotted on the first two principal coordinates for **(A)** hand and hand associated comparisons for all studies combined clustered by Phenotype. Hand versus hand associated PCoA for **(B)** Hospital: Lax et al. 2017, **(C)** air force: Sharma et al. 2019 **(D)** Dorm: Richardson et al. 2019, and **(E)** House: Lax et al. 2014. **(F)** Table of AUC for the unpermuted data in a LOSO design. A red box indicated the p-value was significant for the left out study.

As we might expect, given the weak separation observed between hand and hand-associated samples, our log_10_ p-value-p-value plots had very little agreement between studies, indicated by sparsely populated first and third quadrants **(Fig. S9)**. Additionally, few bacteria were significant across multiple p-value-p-value plots. However, *Sphingomonas,* a soil-associated bacterium, was significantly higher in floor samples for the dorm, hospital, and Air Force studies **(Fig. S9A-C)**.

## Discussion

In this study, we integrated four built environment microbiome datasets using an ontology which grouped samples as hands, hand-associated surfaces, or floors to evaluate reproducibility both within and across studies. Despite substantial differences in environments, sampling strategies, and sequencing protocols, we identified a markedly consistent bacterial signal that consistently distinguished floor samples from those influenced more directly by human contact. The reproducibility of the hand vs floor and hand-associated surfaces vs floor comparisons was evident in PERMANOVA, PCoA, intra-study inference, and leave-one-study-out machine learning analyses, all of which converged on the same broad conclusion: floor microbiomes are strongly and consistently enriched for soil-associated taxa, while hands and hand-associated surfaces carry bacterial profiles shaped by human skin.

Temporal analyses revealed variability within some studies, most notably in one hospital timepoint, but this variation did not obscure the broader cross-study patterns. The weakest and least reproducible contrast was hands and hand-associated surfaces, reflecting the bidirectional bacterial exchange known to occur between human skin and frequently touched objects and the importance of reproducible studies to validate findings of individual studies.

A key implication of our findings is that individual studies can give a misleading picture of built-environment microbiome. For example, the hospital dataset contained one anomalous timepoint (2013/07) in which the communities of hand-associated surfaces and floor were far less distinct than in any other timepoint or study. If this outlier were viewed as representative, one might conclude that hand-associated and floor microbiomes are poorly differentiated or temporally unstable, a conclusion that is strongly contradicted by the remaining timepoints and by external studies. This underscores the risks of inferring general principles of built-environment microbiology from any single study: idiosyncrasies of building type, resident turnover, cleaning routines, sample size, or temporal context can obscure the underlying reproducible patterns that only become clear through integrated, cross-study analysis.

Additionally, differences in cleaning regimens or other unmeasured environmental variables likely contributed to the heterogeneity we observed across studies and timepoints. Built environments differ not only in occupancy but also in disinfectant type, cleaning frequency, ventilation patterns, humidity, and traffic flow which are all correlated with microbiome structure^38,39^. For instance, the anomalous hospital timepoint may reflect an undocumented shift in cleaning practices, staffing, patient turnover, or environmental conditions that altered the balance between human-associated and soil-derived taxa.

When we compared hand-associated surfaces to hands, the study of Air Force dormitories had greater separation between surface types in comparison to the hospital study. We postulate this is the effect of the Air Force dormitories having stable residents across time while the residents of the hospital were more transient, leading to disparate microbiomes across time in the hospital study. Because few built environment studies comprehensively document these contextual variables, it is difficult to disentangle biological signals from building-level interventions. This highlights the need for more standardized metadata reporting, as well as careful interpretation: What appears to be a biological difference within a single study may instead reflect changes in cleaning protocol or other uncontrolled environmental perturbations.

Despite these sources of variation, we were able to identify trends in our longitudinal analyses. Built-environment microbiomes showed substantial cross-study overlap, with samples clustering more strongly by ecological niche (hand, hand-associated surface, floor) than by study of origin. This pattern held even before batch correction and persisted after applying DEBIAS-M, suggesting that the underlying ecological structure of the built environment is sufficiently strong and consistent across buildings and that the signal can overcome study-specific methodological variation. This relative insensitivity to different laboratory workflows enhances the interpretability and reproducibility of built-environment research and suggests that the field may be amenable to cross-study synthesis.

Despite hand and hand-associated surfaces ontological groups typically having larger sample sizes, most of the bacteria which were significantly present across multiple studies tended to be more abundant in floor samples. Other studies have found that floors have higher bacterial abundance than other surfaces^40^. Prior research has also found that shoes act as vectors for outdoor microbiota, even after a considerable number of steps,^7^ which causes a convergence in microbiome between floors and soil.^7,41^ Given that most of our built environment habitats were public or shared spaces (e.g., dorms and hospitals) it was unsurprising that several of the floor bacteria we found are soil bacteria, since even within dorms, inhabitants tend to wear their shoes. It was also unsurprising that our intra-study comparison (PCoA and PERMANOVA) and inter-study comparisons (random forest and p-value-p-value plots) were highly discriminant between the microbiome of hands and floors.

Our results were in line with other non-integrated comparisons between different built environments which found commonalities in the bacteria present in built environment habitats. Phyla *Actinobacteria* (which comprises *Lawsonella and Cutibacterium)*, *Proteobacteria*, *Firmicutes* and *Bacteroidetes* are highly represented in the bacteria present on skin,^42^ and given the populating effect of skin on built environment surfaces, it is unsurprising there is overlap between common skin bacteria and the bacteria of the built environment.^8,10^ Taken together, these results suggest a common microbiome between hand samples which separate them from floor samples Our results also clarify the challenges of batch effects in built-environment meta-analysis. Although the four studies differed in laboratory processing and sequencing, batch correction with DEBIAS-M did not materially improve clustering patterns or classification performance, indicating that biological signal rather than technical noise drove much of the structure observed. Importantly, this suggests that simple ontological grouping can reveal robust ecological signals even without extensive correction.

Although the results presented in this study were compelling, our study was not without limitations. Although the sample size of our individual studies was quite robust (**Table 1**) we were limited in the diversity and number of built environments. Given the differences between non-shared residences such as houses and other built environments, including more residences such as houses and apartments in future would be of interest. Additionally, due to the low sample size of each timepoint, we were not able to include every timepoint for every study leading to an incomplete view of the true variation across time. We also used “inner elbow” to represent the “hand” sample for the Air Force study for the hand versus floor comparison, although this did not appear to be an issue with that study having the highest leave out AUC of 0.973, indicating lack of spatial disparity in that region of the skin microbiome.^43^

Together, these findings demonstrate that there are reproducible patterns of bacterial niche partitioning across diverse built environments, when samples are classified using a consistent ontology. Across study, floors harbor outdoor-derived taxa, hands retain human-skin signatures, and hand-associated surfaces exhibit intermediate profiles. This stable ecological structure, at the genera level, provides a reliable foundation for future research, enabling cross-study comparability amongst 16S rRNA gene studies and offers a baseline against which deviations or environmental interventions, such as changes in cleaning practices or ventilation, can be interpreted. By showing that simple ontological categories produce predicable ecological signals across buildings, populations, and timepoints, our results indicate that built environment microbiomes are not idiosyncratic. As additional datasets from a wider range of environments become available, this framework can be extended to compare ecological signals across ontological niches, strengthen forensic and public-health applications, and advance our understanding of how humans shape the microbial communities of the spaces they inhabit.

## Conclusion

This study shows there is a common microbiome which separates floors from hands and hand associated surfaces. We highlighted the importance of comparing temporal reproducibility using multiple timepoints and studies to correctly characterize microbial changes. Despite batch effects and temporal variation there is a common signal across study which separates hands and hand associated surfaces from floors. Consistently, soil associated taxa are enriched on floors relative to hands and their associated surfaces, while hand associated bacteria are enriched on hands and hand associated surfaces suggesting populating effects. In light of these findings, we highlighted the necessity to assess microbiome findings of multiple studies to gleam correct insights as to the true bacterial composition of built environment structures.

## List of abbreviations

LOSO: leave-one-study-out
PCoA: Principal Coordinate Analysis
PERMANOVA: Permutational Multivariate Analysis of Variance

## Declarations

All manuscripts must contain the following sections under the heading ‘Declarations’:

Ethics approval and consent to participate <- do we need this if we reanalyzed publicly available data?

Consent for publication

## Availability of data and material

The datasets analyzed during the current study are available in the NCBI SRA repository, under accession number PRJEB14474 (https://www.ncbi.nlm.nih.gov/bioproject/?term=PRJEB14474); PRJEB6292 (https://www.ncbi.nlm.nih.gov/bioproject/?term=PRJEB6292); PRJEB33050 (https://www.ncbi.nlm.nih.gov/bioproject/?term=PRJEB33050); PRJEB26708 (https://www.ncbi.nlm.nih.gov/bioproject/?term=PRJEB26708). All scripts used to generate the figures ae available at GitHub (https://github.com/Abeoseh/builtenv_sra)

## Authors’ contributions

A.B.F: manuscript preparation, figure generation, data analysis, conception of ideas

A.A.F: manuscript preparation, conception of ideas

I.B: data analysis

## Acknowledgements

We would like to thank Megan Hill for aiding in the revision of the manuscript.

---

**Table S2:**
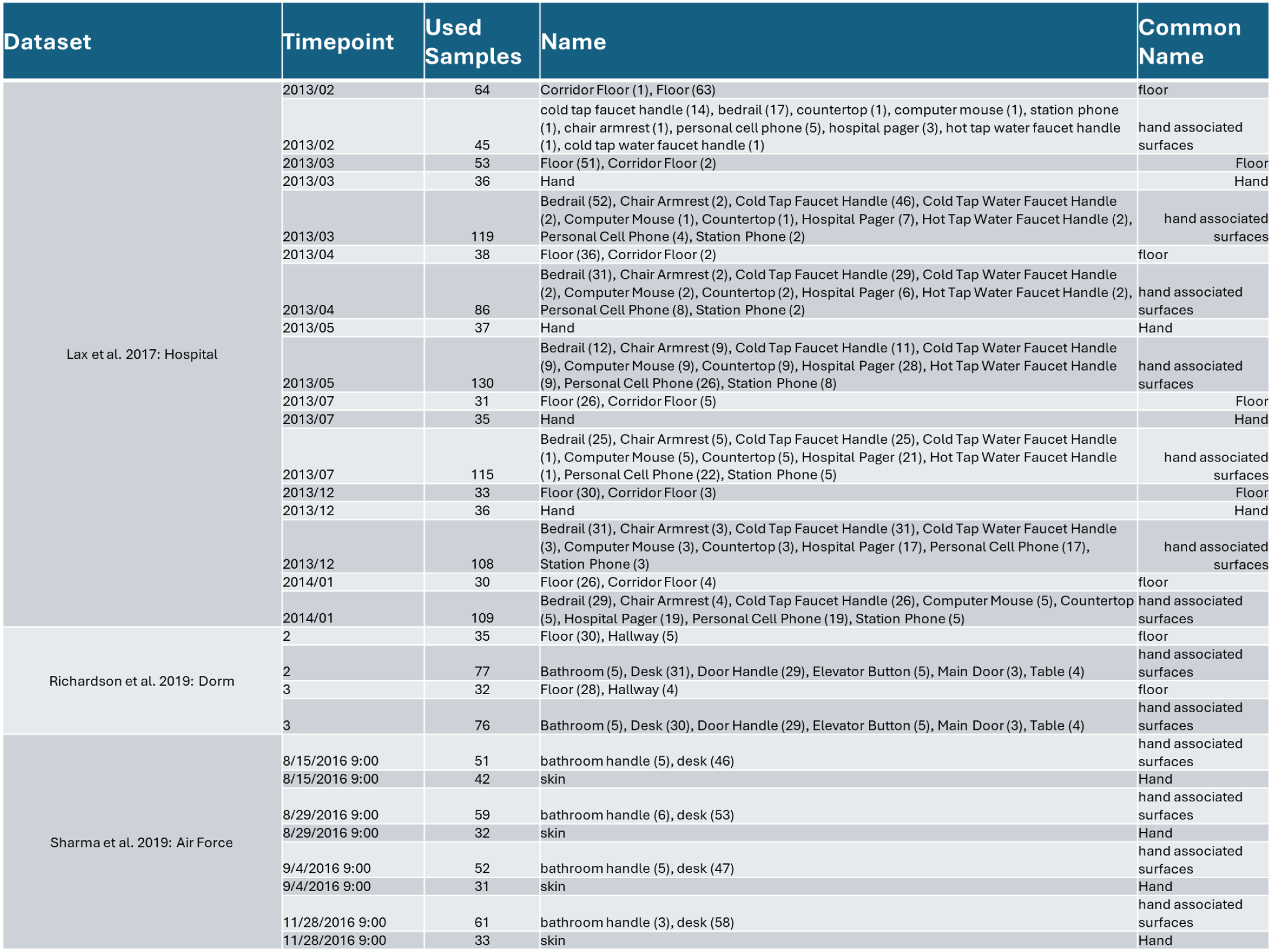
Study characteristics stratified by timepoint. The amount used (Samples), the samples used in the study (Name), and the common name we used in our analysis (Common Name) for the hand, hand associated surfaces, and floor samples for each timepoint in each study (Dataset). Sharma et al. 2019: Air Force swabbed the inner elbow.

**Figure S10:**
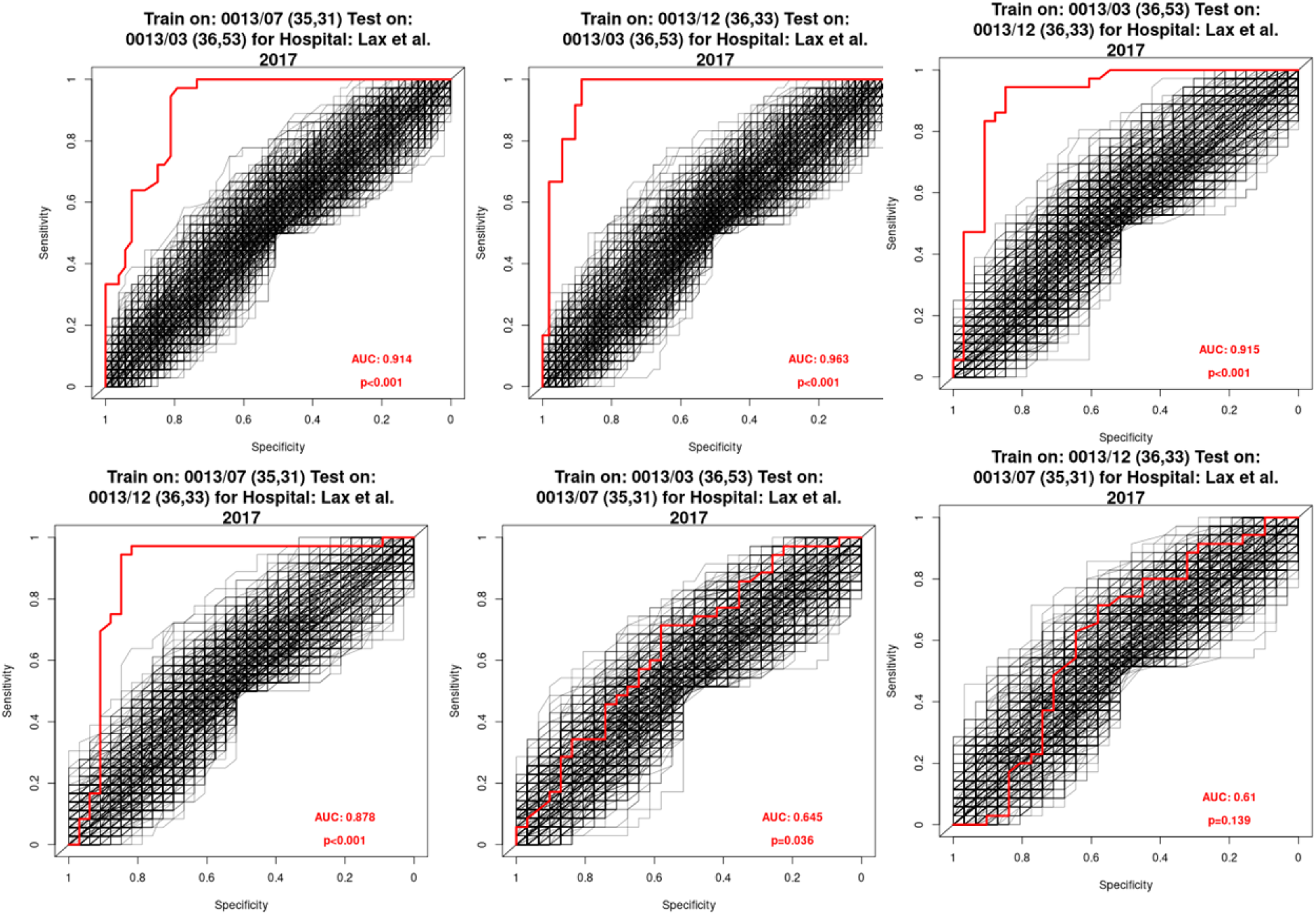
ROC curves after differentiating the microbiome of hands and floors across time in the Hospital study. ROC curves for the pairwise timepoint comparisons, when testing and training on different timepoints. The bootstrap p-values is the proportion in which the AUC of the non-permuted data was equal to or greater than the AUC of the permuted data.

**Figure S11:**
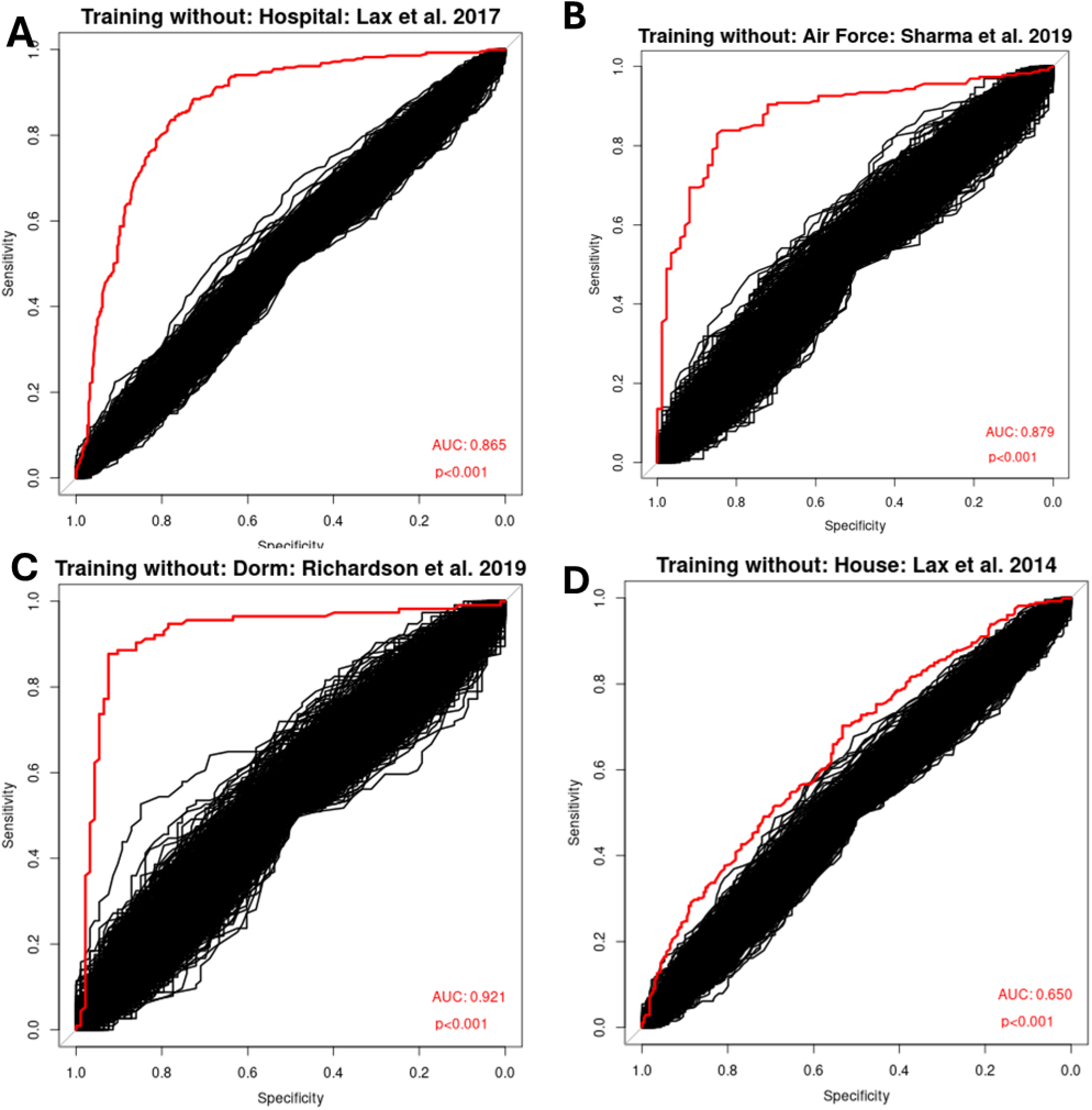
ROC curves of the un-permutated and permutated data after comparing the microbiome of hands and floors across study. pre-DEBIAS-M hand and floor leave one study out **(A-D)** ROC curves of the un-permutated data (red) and 100 permutations (black) for the **(A)** hospital, **(B)** air force, **(C)** dorm, and **(D)** house studies. The AUC is for the un-permutated data and the p-value is number of times the AUC of the permutations were equal to or greater than the AUC of the actual data.

**Figure S12:**
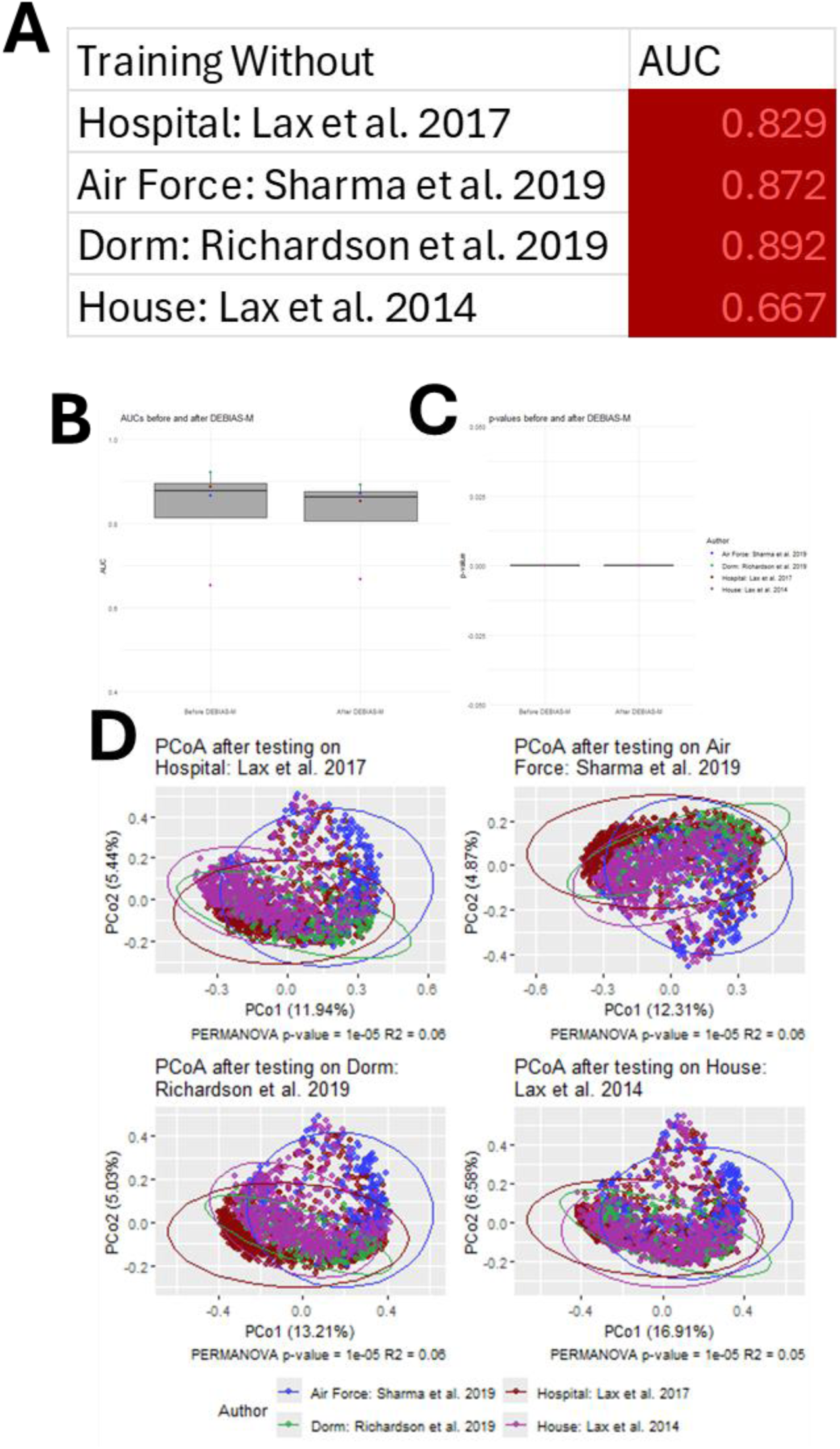
Across study microbiome comparison of hands and floors after batch correction. t-test between **(A)** AUCs and **(B)** p-values before and after DEBIAS-M for hand and floors. **(C)** Intra-study PERMANOVA and PCoA between studies with 100,000 permutations. **(D)** PCoA and PERMANOVA of the data post DEBIAS-M.

**Figure S13:**
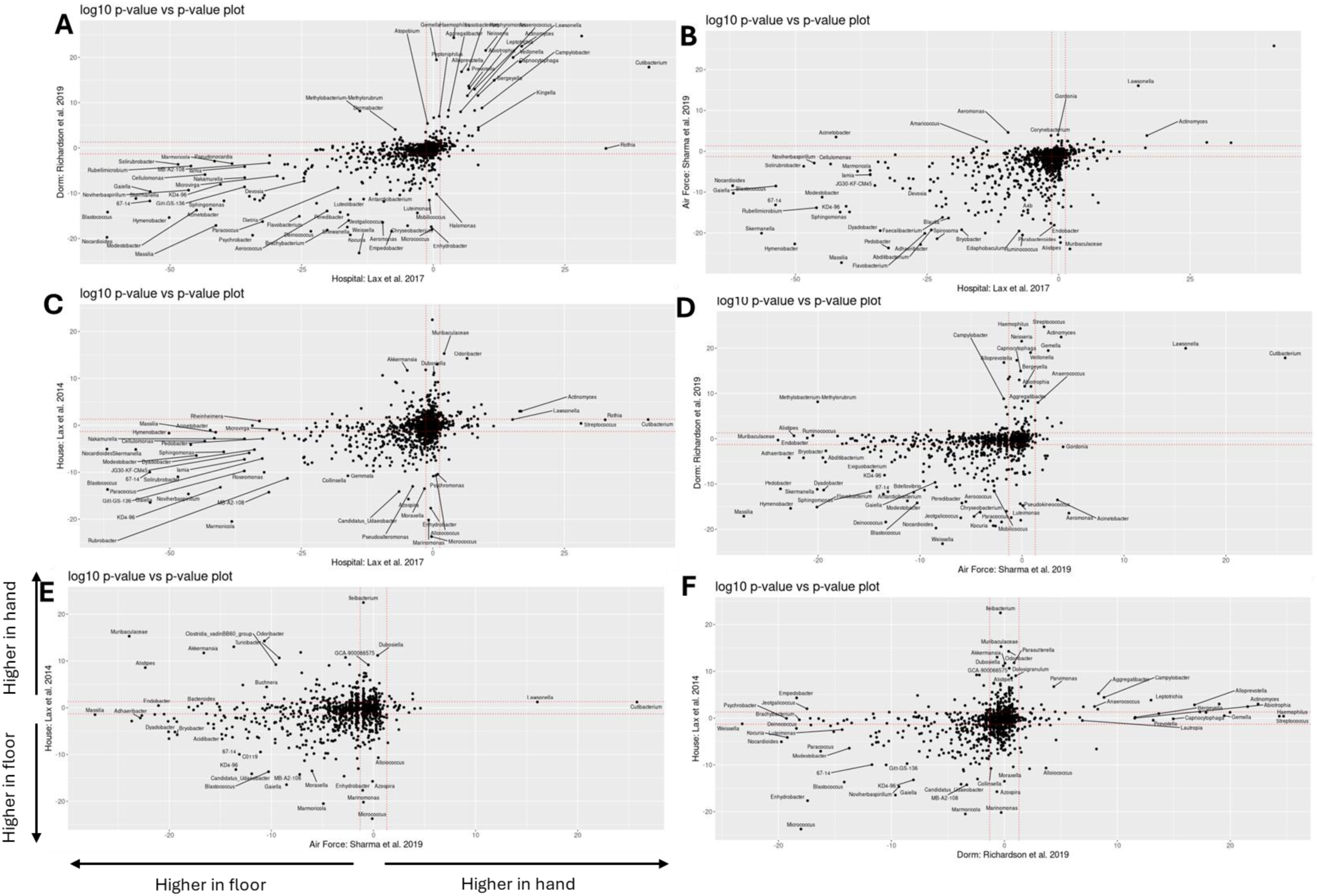
Agreement and Disagreement between studies of the bacteria which separate hands from floors. **(A-F)** p-value-p-value plots for hand and floor samples within studies where quadrant 1 shows agreement between studies that a given bacterium is significantly higher in hand samples, and quadrant 3 shows inter-study agreement that a given bacterium is significantly higher is floor samples. Red lines are the demarcation for the log10 p-value cutoff of 0.05.

**Figure S14:**
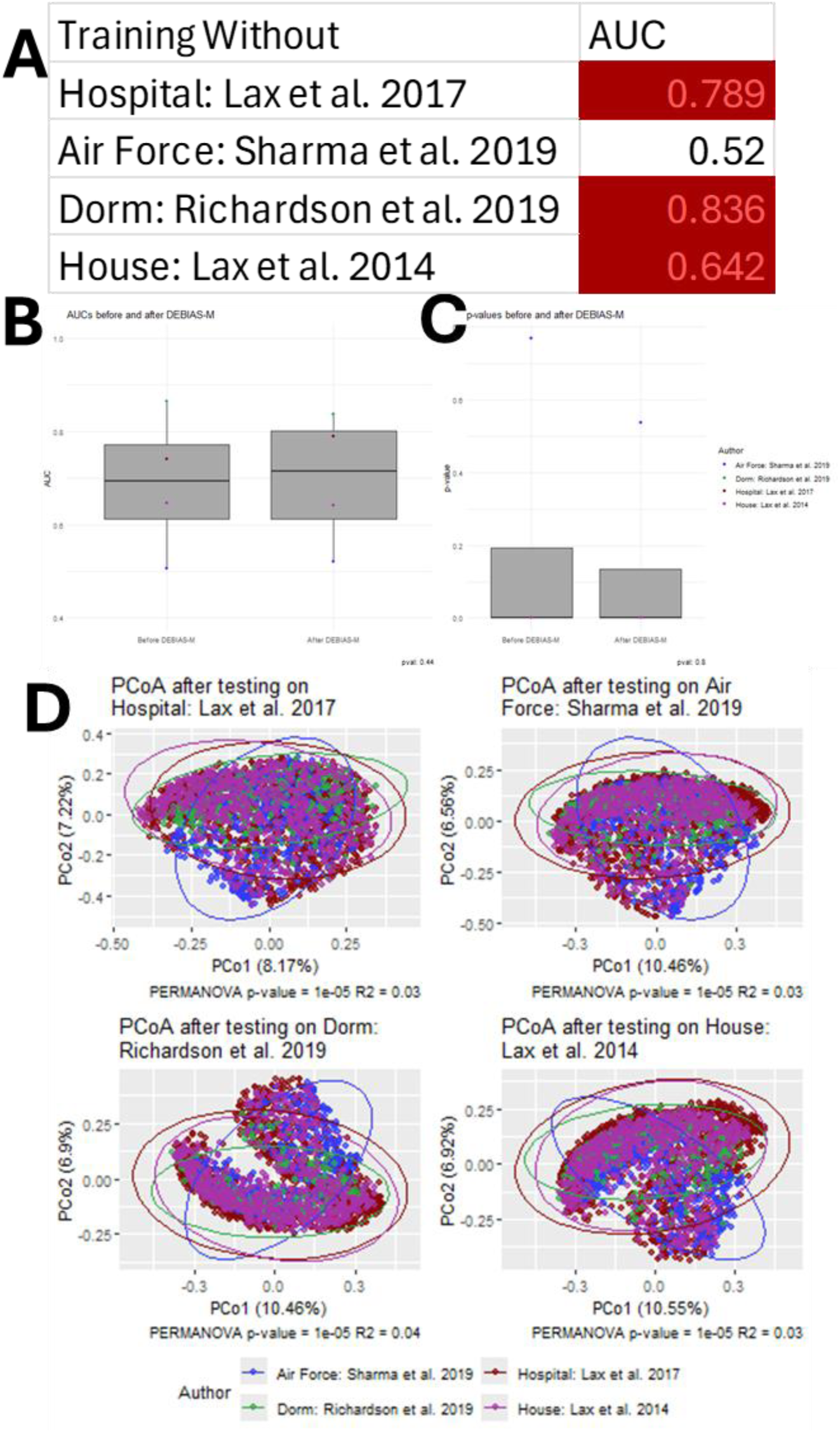
Across study microbiome comparison of hand associated surfaces and floors after batch correction. t-test between **(A)** AUCs and **(B)** p-values before and after DEBIAS-M for hand associated surfaces and floors. **(C)** Intra-study PERMANOVA and PCoA between studies with 100,000 permutations. **(D)** PCoA and PERMANOVA of the data post DEBIAS-M.

**Figure S15:**
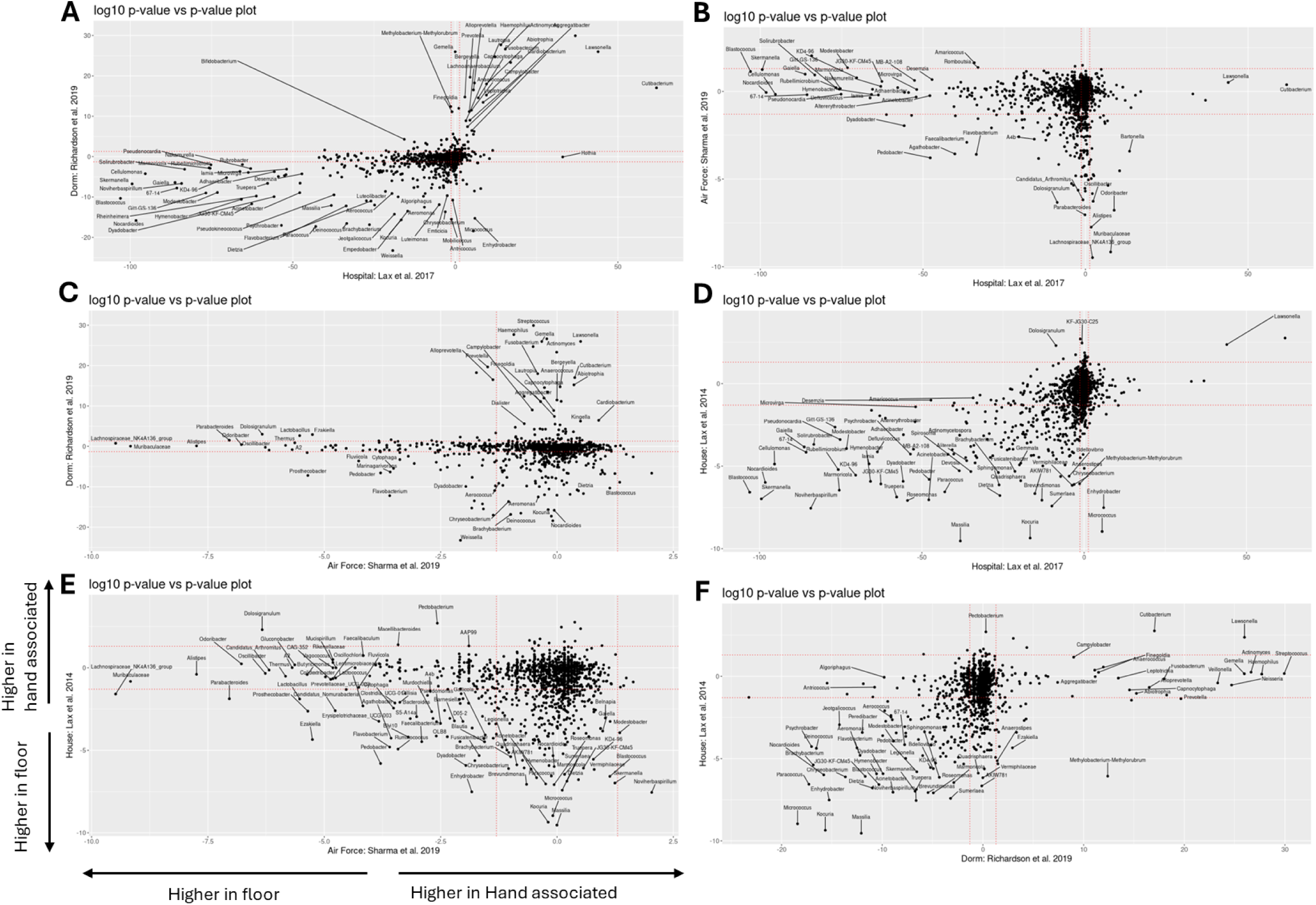
Agreement and Disagreement between studies of the bacteria which separate hand associated surfaces from floors. **(A-F)** p-value-p-value plots for hand associated and floor samples within studies where quadrant 1 shows agreement between studies that a given bacterium is significantly higher in hand associated samples, and quadrant 3 shows agreement between studies that a given bacterium is significantly higher is floor associated samples. Red lines are the demarcation for the log10 p-value cutoff of 0.05.

**Figure S16:**
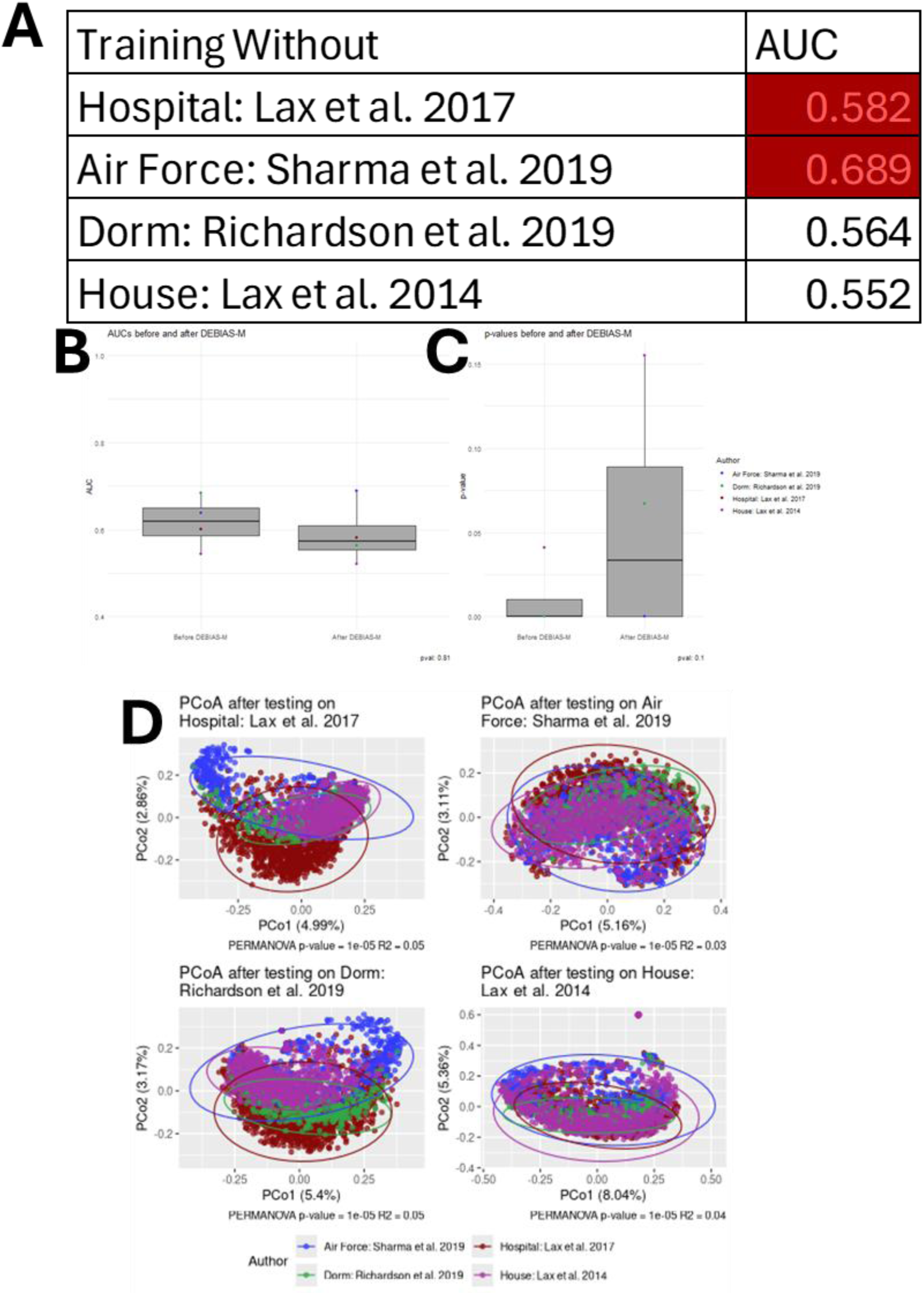
Across study microbiome comparison of hands and hand associated surfaces after batch correction. t-test between **(A)** AUCs and **(B)** p-values before and after DEBIAS-M for hand and hand associated surfaces. **(C)** Intra-study PERMANOVA and PCoA between studies with 100,000 permutations. **(D)** PCoA and PERMANOVA of the data post DEBIAS-M.

**Figure S17:**
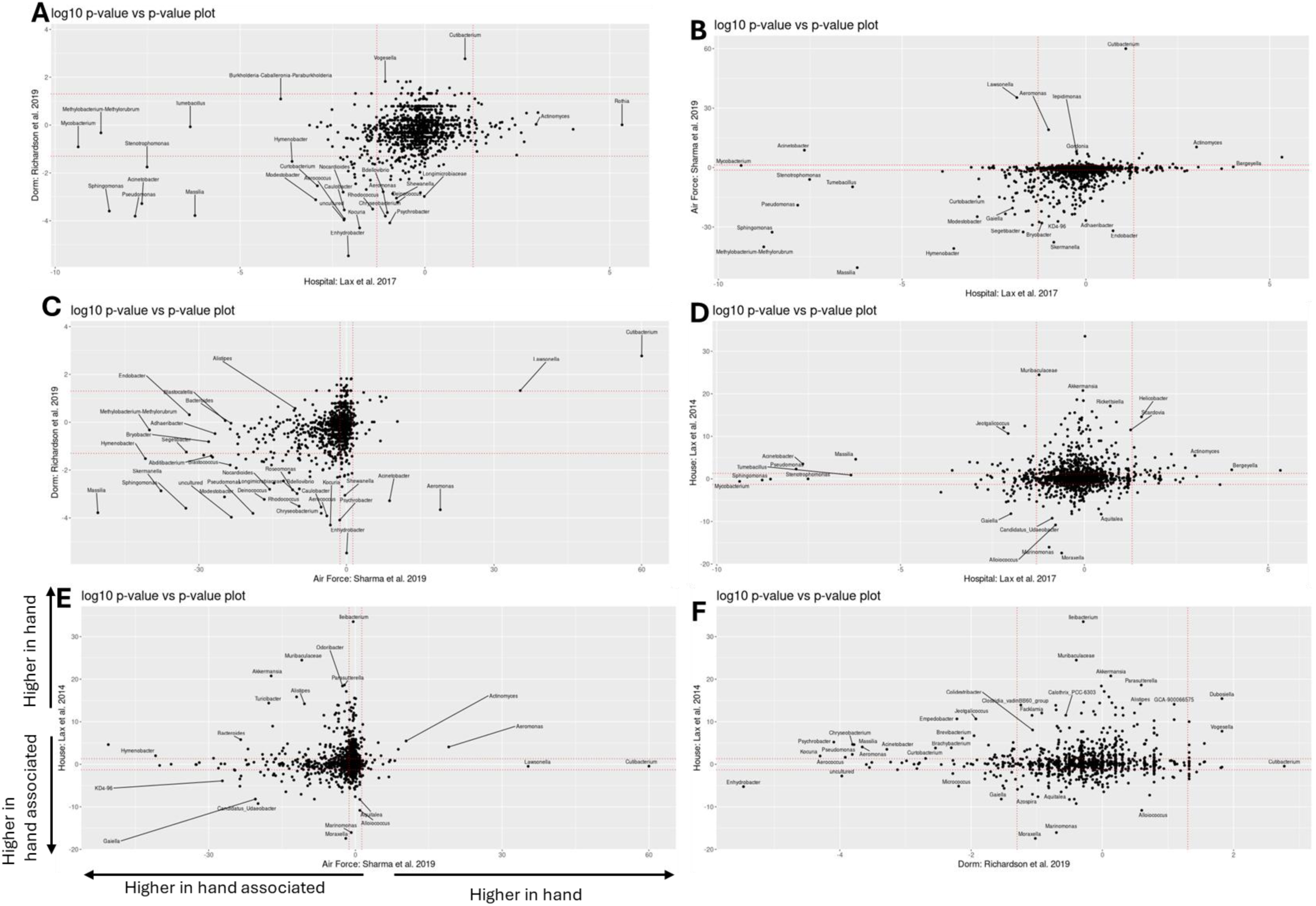
Agreement and Disagreement between studies of the bacteria which separate hands from hand associated surfaces. **(A-F)** p-value-p-value plots for hand and hand associated samples within studies where quadrant 1 shows agreement between studies that a given bacterium is significantly higher in hand samples, and quadrant 3 shows agreement between studies that a given bacterium is significantly higher is hand associated samples. Red lines are the demarcation for the log10 p-value cutoff of 0.05.

